# Efficient multiple gene knock-out in *Colletotrichum higginsianum* via CRISPR-Cas9 ribonucleoprotein and *URA3*-based marker recycling

**DOI:** 10.1101/2023.04.20.537420

**Authors:** Katsuma Yonehara, Naoyoshi Kumakura, Takayuki Motoyama, Nobuaki Ishihama, Jean-Félix Dallery, Richard O’Connell, Ken Shirasu

## Abstract

Colletotrichum higginsianum is a hemibiotrophic pathogen that causes anthracnose disease on crucifer hosts, including Arabidopsis thaliana. Despite the availability of genomic and transcriptomic information and the ability to transform both organisms, identifying C. higginsianum genes involved in virulence has been challenging due to their recalcitrance to gene targeting and redundancy of virulence factors. To overcome these obstacles, we developed an efficient method for multiple gene disruption in C. higginsianum by combining CRISPR-Cas9 and URA3-based marker recycling systems. Our method significantly increased the efficiency of gene knock-out via homologous recombination by introducing genomic DNA double-strand breaks. We demonstrated the applicability of the URA3-based marker recycling system for multiple gene targeting in the same strain. Using our technology, we successfully targeted two melanin biosynthetic genes, SCD1 and PKS1, which resulted in deficiency in melanisation and pathogenicity in the mutants. Our findings demonstrate the effectiveness of our developments in analysing virulence factors in C. higginsianum, thus accelerating research on plant-fungus interactions.

## 1 INTRODUCTION

Phytopathogenic fungi cause significant crop losses annually, leading to serious economic consequences (Fisher et al., 2012; Oerke, 2006). Among them*, Colletotrichum* spp. are in the top 10 most economically and scientifically important phytopathogenic fungi, due to their wide host range and hemibiotrophic lifestyle (Dean et al., 2012). Members of the genus *Colletotrichum* are responsible for anthracnose disease in thousands of plant species and undergo, in general, two distinct phases during the infection process, namely the biotrophic and necrotrophic phases (Cannon et al., 2012; Münch et al., 2008; Perfect et al., 1999). The infection begins with the development of melanised appressoria from fungal spores, which generate high turgor pressure to penetrate the host plant’s cuticle and cell wall. In the subsequent biotrophic phase, the fungi extend their mycelia within the living host cells to obtain nutrients. The infection is completed in the necrotrophic phase, marked by host cell death and the formation of spores (Cannon et al., 2012). This dynamic lifestyle transition makes *Colletotrichum* spp. valuable subjects for research on plant-fungus interactions.

Within the genus, *Colletotrichum higginsianum* is a suitable model for exploring plant-fungus interactions due to its ability to infect cruciferous plants including *Arabidopsis thaliana*, a species with extensive genomic, transcriptomic resources, and genetic tools (Narusaka et al., 2004; O’Connell et al., 2004). Studies involving genetic crosses of ecotypes with varying resistance to *C. higginsianum* have led to the identification of resistance genes in *A. thaliana* (Birker et al., 2009; Narusaka et al., 2009). On the fungal side, genomic and transcriptomic analyses of *C. higginsianum* have highlighted the potential contribution of secondary metabolites and secreted proteins to infection (Dallery et al., 2017; Kleemann et al., 2012; O’Connell et al., 2012; Tsushima et al., 2019; Tsushima and Shirasu, 2022). For instance, the secondary metabolite higginsianin B and extracellular LysM proteins of *C. higginsianum* have been found to suppress plant immune responses (Dallery et al., 2020; Takahara et al., 2016). Further, targeted gene disruption is available in *C. higginsianum* through *Agrobacterium tumefaciens*-mediated or PEG-mediated protoplast transformation, allowing the search for pathogenicity genes (Huser et al., 2009; Ushimaru et al., 2010), and homologous recombination utilising donor DNA enables disruption of a target gene. However, the molecular mechanisms underlying the infection of *C. higginsianum* are not yet fully understood, probably due to the recalcitrance of some genes to manipulation (Huser et al., 2009; Ushimaru et al., 2010) and the redundancy of pathogenicity genes (Collemare et al., 2019; Kleemann et al., 2008). Therefore, the development of high-efficiency multiple gene disruption in *C. higginsianum* would be valuable for uncovering plant-fungus interactions, but such technology is currently unavailable for this species.

The clustered regularly interspaced short palindromic repeats (CRISPR)-associated protein 9 (Cas9) system is a revolutionary tool for genome editing, increasing the efficiency of gene disruption via homologous recombination in various organisms. It operates by inducing double strand breaks (DSBs) in genomic DNA (Belhaj et al., 2015; Sander and Joung, 2014; Schuster and Kahmann, 2019) using ribonucleoproteins (RNPs) comprising Cas9 proteins and guide RNAs that target specific sequences. The resulting repair of DSBs through non-homologous end joining (NHEJ) or homology-directed repair (HDR) pathways (Capecchi, 1989; Pastink et al., 2001) enables targeted gene manipulation. Artificially inducing DSBs in target regions with CRISPR-Cas9 significantly increases the efficiency of homologous recombination using donor DNA (Devkota, 2018; Gratz et al., 2014). This technique has been demonstrated and utilised in several phytopathogenic fungi, including *Magnaporthe oryzae* (Arazoe et al., 2015; Foster et al., 2018), *Botrytis cinerea* (Leisen et al., 2020), *Penicillium expansum* (Clemmensen et al., 2022), *Colletotrichum sansevieriae* (Nakamura et al., 2019) and *Colletotrichum orbiculare* (Chen et al., 2023; Yamada et al., 2023). Nevertheless, despite its effectiveness, CRISPR-Cas9 has not yet been applied to *C. higginsianum*.

A selection marker recycling system in combination with homologous recombination has enabled the efficient disruption of multiple genes in a variety of fungi. The *URA3*-based marker recycling system that relies on the versatile *URA3* gene encoding orotidine-5’phosphate decarboxylase involved in uridine synthesis (Weld et al., 2006) has been widely used in fungi, such as yeast (Alani et al., 1987), *Aspergillus* (d’Enfert, 1996; Nielsen et al., 2006; Oakley et al., 1987), and *C. orbiculare* (Kumakura et al., 2019). In this system, using *ura3* mutants exhibiting uridine auxotrophy as recipients, strains can be selected, in which the target loci in the genome are replaced with donor DNA containing an *URA3* expression cassette, on selection media without uridine. Negative selection can then be used to select for strains that lost the *URA3* cassette using 5-fluoroorotic acid (5-FOA), an analogue of uridine precursor, because 5-FOA is converted into the toxic compound fluorouracil by URA3 (Boeke et al., 1984; Flynn and Reece, 1999). The *URA3* cassette can be removed from the genome via homologous recombination if it is flanked by two homologous sequences. This selection system can be repeated as many times as necessary to achieve multiple gene disruption with one selection marker. The *URA3*-based marker recycling system should be applicable to *C. higginsianum*, as fungi generally possess the *URA3* gene.

Here, we report the establishment of an efficient multiple gene disruption method in *C. higginsianum* combining CRISPR-Cas9 and *URA3*-based marker recycling. By using *in vitro* formed CRISPR-Cas9 RNPs in the PEG-mediated transformation of *C. higginsianum* protoplast with donor DNAs, we observed a significant increase in gene knock-out efficiency. Using *URA3*-based marker recycling, we demonstrated the capacity for multiple gene disruption in the same strain by sequentially targeting two melanin biosynthesis genes, *SCD1* and *PKS1*. The *scd1* and *pks1 scd1* mutants exhibited melanin deficiency and loss of pathogenicity on *A. thaliana* leaves, demonstrating the effectiveness of this method for phenotypic analysis of virulence-related genes. Our results establish a highly efficient multiple gene disruption method in *C. higginsianum*, which has the potential to aid the identification of previously unidentified virulence factors.

## 2 RESULTS

### 2.1 CRISPR-Cas9 RNP enhances the rate of donor-DNA mediated HDR in *C. higginsianum*

To identify the *URA3* homologue in the *C. higginsianum* genome, we employed the NCBI BlastP tool with default settings and utilised *S. cerevisiae* URA3p (GenBank ID: NP_010893.3) as the query sequence. As previously reported (Kumakura et al., 2019), *URA3* homologues are generally well-conserved among phytopathogenic fungi. Accordingly, we identified one *URA3* homologue in the *C. higginsianum* genome (Locus tag: CH63R_12904) (Figure S1). Protoplast transformation-mediated attempts to disrupt *URA3* were unsuccessful, leading us to introduce CRISPR-Cas9 technology to induce DSBs and promote homologous recombination with donor DNA. The transformation method with CRISPR-Cas9 RNPs consists of three steps (Figure 1a). First, Cas9 recombinant proteins and synthetic gRNAs are mixed *in vitro* to form RNPs. Next, *C. higginsianum* protoplasts are transformed using donor DNA and CRISPR-Cas9 RNPs. Finally, selective media and PCR-based screening techniques are employed to identify strains with the intended HDRs.

**Figure 1.**
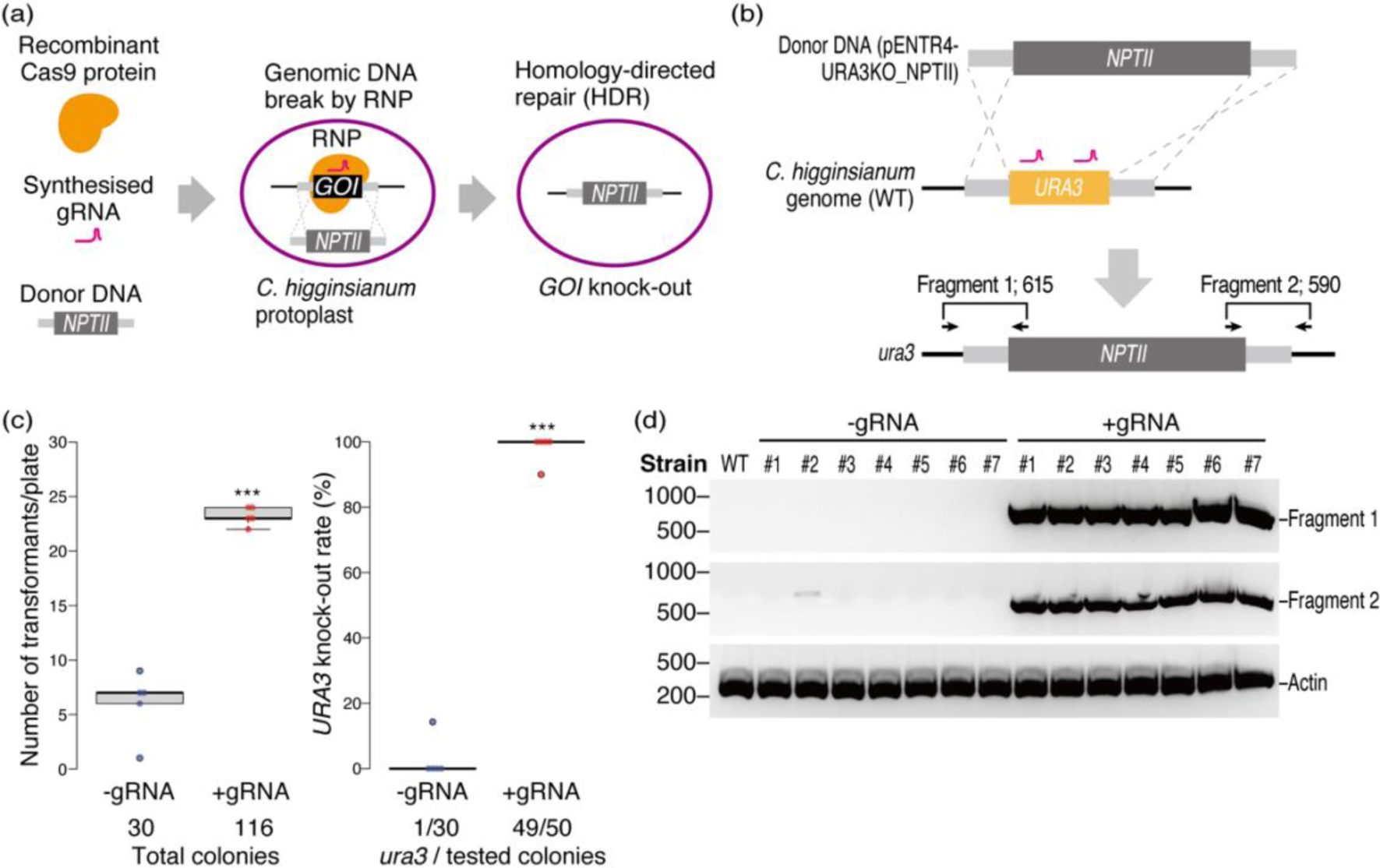
CRISPR-Cas9 RNP promotes homologous recombination with donor DNA in *C. higginsianum*. (a) Schematics of CRISPR-Cas9 RNP-mediated HDR. Firstly, recombinant Cas9 proteins (in orange) and synthesised gRNAs (in magenta) targeting the gene of interest (GOI) were mixed *in vitro* to form RNPs. Secondly, *C. higginsianum* protoplasts were transformed with RNPs and donor DNAs that have the selection marker, *NPTII*, flanked by two homology arms. Finally, the strain in which the GOI was replaced with the selection cassette was isolated by combining selection media and PCR-based screening. (b) Construction design of *URA3* knock-out. The donor DNA has a selection marker, the *NPTII* expression cassette, flanked by 0.5 kb homology arms depicted as light gray boxes. Arrows represents the primers to amplify “Fragment 1” and “Fragment 2” existing specifically in the genome of *ura3*. (c) Number of transformants and *URA3* knock-out rate. The left panel shows the number of transformants per plate, and the right panel shows the *URA3* knock-out rate per plate, respectively (*n*=5). The “-gRNA” and “+gRNA” represents the results without and with gRNAs, respectively. The asterisks indicate statistical difference (*p* < 0.001, Welch’s t test). The knockout of *URA3* was evaluated by PCR screening as shown in (d). (d) PCR screening of *ura3* mutants. PCRs were performed using each colony on MA containing 500 µg/ml G418 with primer sets depicted in (b). Results from randomly selected seven colonies from −gRNA and +gRNA transformants are shown. The 238 bp fragment of the *C. higginsianum Actin* gene (CH63R_04240) was designated as Actin. The numbers on the left of the gels indicate the position of the DNA size marker (bp).

The donor DNA for *URA3* knock-out was designed to have a *neomycin phosphotransferase II* (*NPTII*) expression cassette conferring G418/geneticin resistance, flanked with two homology arms that matched upstream and downstream regions of the *URA3* coding sequence (CDS) (Figure 1b). Two gRNAs were designed to target the CDS of *URA3*. Wild-type *C. higginsianum* protoplasts were transformed using CRISPR-Cas9 RNPs and the donor DNA. A negative control was performed without gRNA. Our result showed that CRISPR-Cas9 RNPs increased the number of colonies by approximately four-fold compared to the negative control (Figure 1c, left panel). We then screened the genomic DNA of colonies by PCR to determine the rate of *URA3* knock-out. We designed two primer sets to amplify two different fragments exclusively present in the genomic DNA from *ura3* mutants that underwent double cross-over with the donor DNA (Figure 1b). PCR screening with these primer sets demonstrated that CRISPR-Cas9 RNPs greatly increased the HDR rate (Figure 1c, right panel, d). The *ura3* rate was 98% with gRNA (+gRNA), whereas it was only 3.3% without gRNA (-gRNA). Hence, we successfully generated *ura3* mutants that can be used to establish a marker recycling system.

### 2.2 *ura3* mutants show uridine auxotrophy and 5-FOA insensitivity

To confirm the uridine auxotrophy of *ura3* mutants, we assessed the colony diameters of *C. higginsianum* strains cultured on media with or without uridine. As expected, the *ura3* mutants exhibited significantly smaller colony diameters compared to the wild type, while the phenotype was restored upon uridine supplementation, providing evidence of their uridine auxotrophy (Figure 2a, b). To investigate the 5-FOA sensitivity of *C. higginsianum*, we evaluated its impact on colony growth. Fungal URA3, in general, is involved in the conversion of 5-FOA to toxic compounds (Boeke et al., 1984; Rossmann et al., 2011). The addition of 5-FOA inhibited the growth of wild-type *C. higginsianum* colonies, indicating their sensitivity to 5-FOA (Figure 2a, b). In contrast, *ura3* mutants were resistant to 5-FOA supplementation, thereby confirming the 5-FOA sensitivity of *C. higginsianum* via *URA3* gene. Therefore, the uridine auxotrophy and the 5-FOA insensitivity of *ura3* mutants indicate the loss of URA3-associated enzymatic activities.

**Figure 2.**
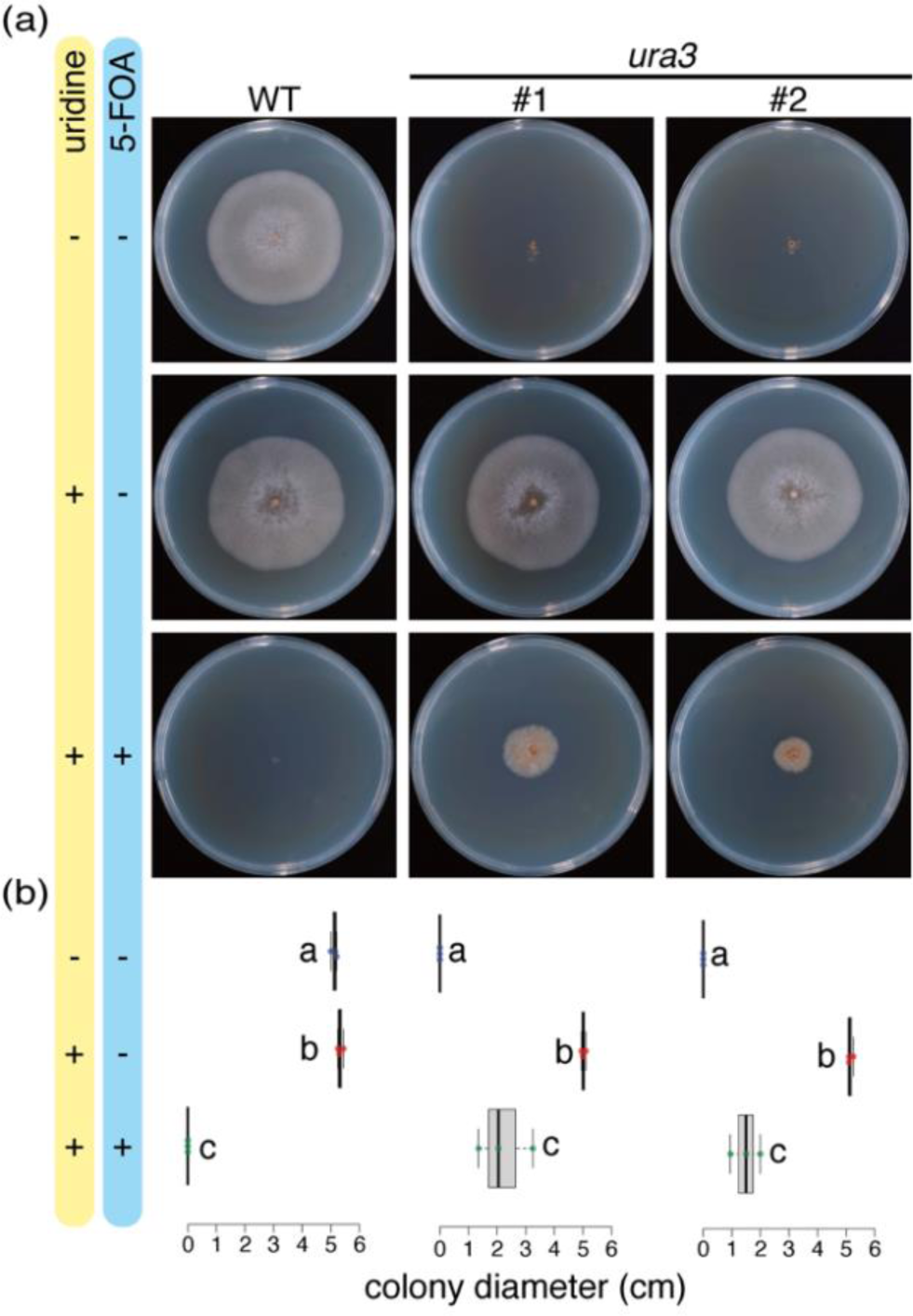
*ura3* mutants show uridine auxotrophy and 5-FOA insensitivity. (a) Colonies of wild-type *C. higginsianum* and two independent *ura3* mutants on MA. Photographs were taken at seven days post-culture. The “+” and “-” indicate the presence or absence of uridine or 5-FOA, respectively. (b) Box plots of colony diameters of *C. higginsianum* strains shown in (a). Different characters in each graph indicate statistical significance (Tukey HSD, *n*=3).

### 2.3 *URA3*-based marker recycling system is applicable in *C. higginsianum*

To evaluate the feasibility of using the *URA3* expression cassette as a selection marker in *C. higginsianum*, we aimed to target the *SCD1* gene that encodes a scytalone dehydratase involved in the synthesis of 1,8-DHN-melanin (Kubo et al., 1996). Mutations in *SCD1* are predicted to result in melanin deficiency and scytalone accumulation. By performing a BlastP search using *C. orbiculare* SCD1 (GenBank ID: TDZ18313.1) (Kubo et al., 1996) as a query, we identified a homologue encoded by the *C. higginsianum* genome (Locus tag: CH63R_00358). To knock out the *C. higginsianum SCD1* gene in the *ura3* mutant background, we employed a combination of the CRISPR-Cas9 system and the donor DNA-mediated HDR (Figure 2b). The donor DNA contained a constitutive *URA3* expression cassette (*TEF-URA3*) flanked by homology arms (Figure 3a). We designed two gRNAs to target the *SCD1* CDS. PCR-based screening revealed successful gene knock-out of *SCD1*, resulting in strains named *scd1::TEF-URA3 ura3* (Figure 3b, c). Although the number of colonies obtained was not significantly affected by CRISPR-Cas9 RNPs (Figure 3b, left panel), the HDR efficiency increased significantly (Figure 3b, right panel, c). Our findings suggest that the *TEF-URA3* expression cassette can serve as an effective selection marker for transforming *ura3* through the combination with CRISPR-Cas9.

**Figure 3.**
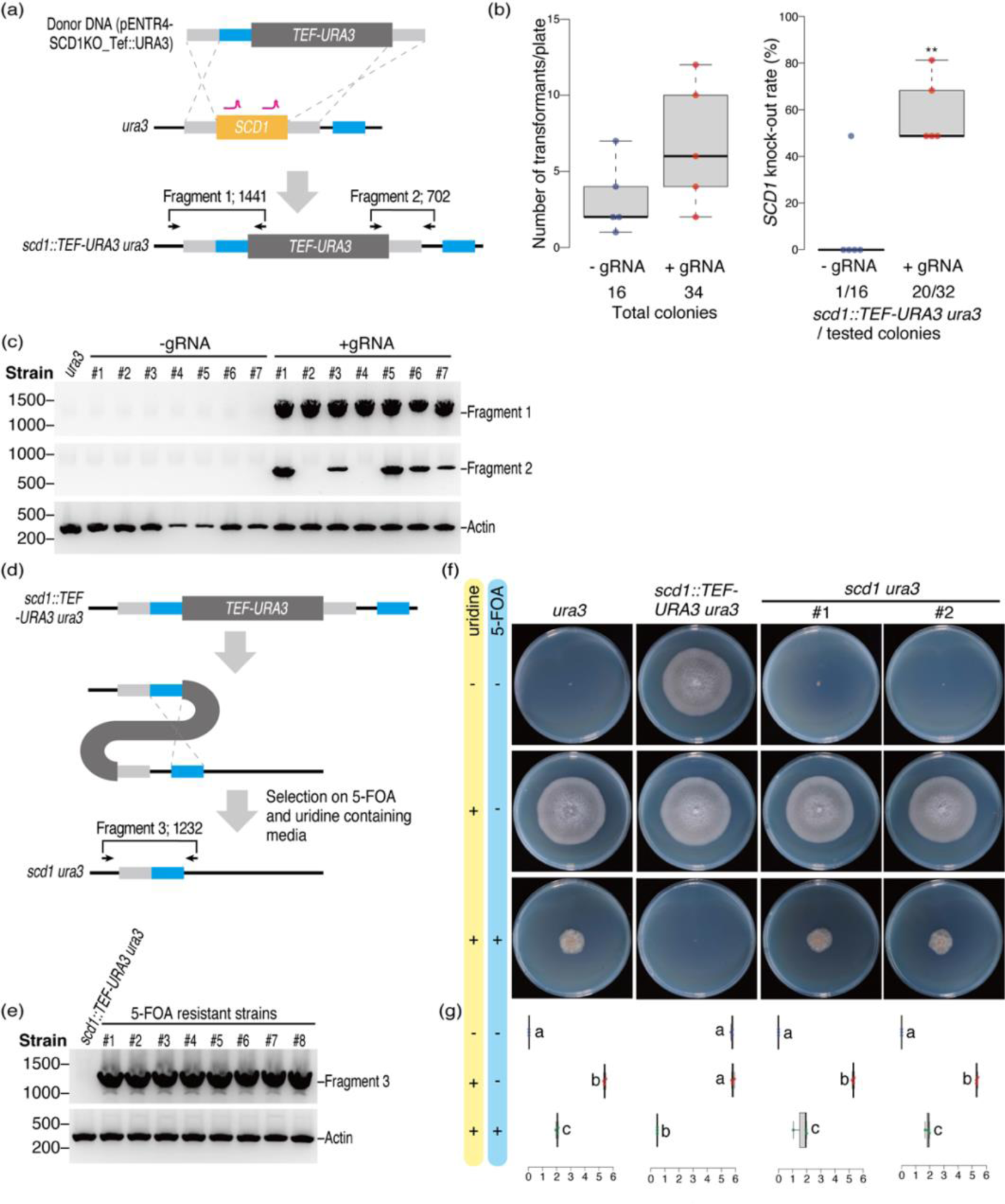
*TEF-URA3* is applicable as both positive and negative selection markers. (a) Construction design of *SCD1* knock-out in *ura3* mutants. The donor DNA contains *TEF-URA3* flanked by 0.5 kb homology arms depicted as light gray boxes. Lines in magenta represent the sites targeted by gRNAs. Light blue boxes represent homology arms to remove *TEF-URA3* from the genome of *scd1::TEF-URA3 ura3* via homologous recombination as shown in (d). Arrows represent primers to amplify “Fragment 1” and “Fragment 2” existing specifically in the genome of *scd1::TEF-URA3 ura3*. (b) Number of transformants and *SCD1* knock-out rate. The left panel shows the number of transformants per plate, and the right panel shows the *SCD1* knock-out rate per plate, respectively (*n*=5). Asterisks indicate statistically significant difference (*p* < 0.01, Welch’s t test). The knockout of *SCD1* was evaluated by PCR screening, as shown in (c). (c) PCR screening of *scd1::TEF-URA3 ura3* mutants. PCRs were performed using primer sets depicted in (a). Results from randomly selected seven colonies from −gRNA and +gRNA transformants are shown. (d) Construction design to remove *TEF-URA3* from *scd1::TEF-URA3 ura3* mutants. Arrows represent primers to amplify “Fragment 3” specifically existing in the genome of *scd1 ura3*. (e) PCR screening of *scd1 ura3* mutants. PCRs were performed using each colony grown on MA containing 10 mM uridine and 1 mg/ml 5-FOA with primer sets depicted in (d). Results of randomly selected eight colonies from transformants are shown. (f) Colonies of *ura3*, *scd1::TEF-URA3 ura3* and two independent *scd1 ura3* mutants grown on MA. Photographs were taken at seven days post-culture. The “+” and “-” indicate the presence or absence of uridine or 5-FOA, respectively. (g) Box plots of colony diameters of *C. higginsianum* strains shown in (f). Different letters in each graph indicate statistical significance (Tukey HSD, *n*=3).

To reuse *TEF-URA3* as a selection marker for sequential gene knock-outs, we next aimed to select strains that had lost *TEF-URA3* from the *scd1::TEF-URA3 ura3* strains. As *TEF-URA3* is flanked by two identical 500-bp sequences (Figure 3d), strains that lost the cassette via homologous recombination could be selected by incubation on media containing 5-FOA and uridine. As expected, *scd1::TEF-URA3 ura3* mutants displayed 5-FOA sensitivity (Figure 3f, middle columns), demonstrating that *TEF-URA3* can be effectively used for negative selection on media containing 5-FOA. To isolate *scd1 ura3* mutants, conidia of *scd1::TEF-URA3 ura3* were cultured on media with 5-FOA and uridine. After several days of incubation, a number of colonies appeared, and genomic DNA PCR analysis revealed the absence of *TEF-URA3* in eight randomly selected colonies. The designated primer amplified 1232-bp fragments in the absence of *TEF-URA3* (Figure 3d), and as expected, the bands were observed from only *scd1 ura3* mutants and not from the parental strain (Figure 3e). Consistent with the loss of *TEF-URA3*, insensitivity to 5-FOA and uridine auxotrophy of *scd1 ura3* were confirmed (Figure 3f, right columns). Thus, our results show that the removal of *TEF-URA3* from *scd1::TEF-URA3 ura3* can be achieved through selection after 5-FOA treatment, enabling the reuse of *TEF-URA3* as a selection marker for sequential gene knock-outs. These results confirm that the *URA3*-based marker recycling system is applicable in *C. higginsianum*.

### 2.3 The *URA3* expression cassette is a reusable selection marker for sequential gene knock-out in *C. higginsianum*

To confirm the reusability of *TEF-URA3* as a selection marker for sequential gene knock-outs in *C. higginsianum*, we targeted *PKS1* in *scd1 ura3* mutants*. PKS1* encodes a polyketide synthase involved in the production of scytalone in the melanin biosynthesis pathway (Takano et al., 1997) (Figure 4a). We identified one *PKS1* homologue in the *C. higginsianum* genome, previously named *ChPKS1* (Locus tag: AB508803) and also referred as *ChPKS19* (Locus tag: CH63R_08918) (Dallery et al., 2017; Ushimaru et al., 2010), by a BlastP search using *C. orbiculare* PKS1 (GenBank ID: TDZ26149.1) as the query. *PKS1* was knocked out in *scd1 ura3* using the same method employed for the *SCD1* knock-out (Figure 4b). PCR-based screening of 10 randomly obtained colonies resulted in five that displayed successful knock-out of *PKS1.* These strains were named *pks1::TEF-URA3 scd1 ura3* (Figure 4c). We then selected strains that had lost *TEF-URA3* from the *pks1::TEF-URA3 scd1 ura3* mutants by culturing them on media containing 5-FOA and uridine. Using PCR-based screening (Figure 4d), we were able to isolate *pks1 scd1 ura3* mutants. Thus, our results demonstrate that the *URA3*-based marker recycling system is applicable to sequential gene knockouts in *C. higginsianum*.

**Figure 4.**
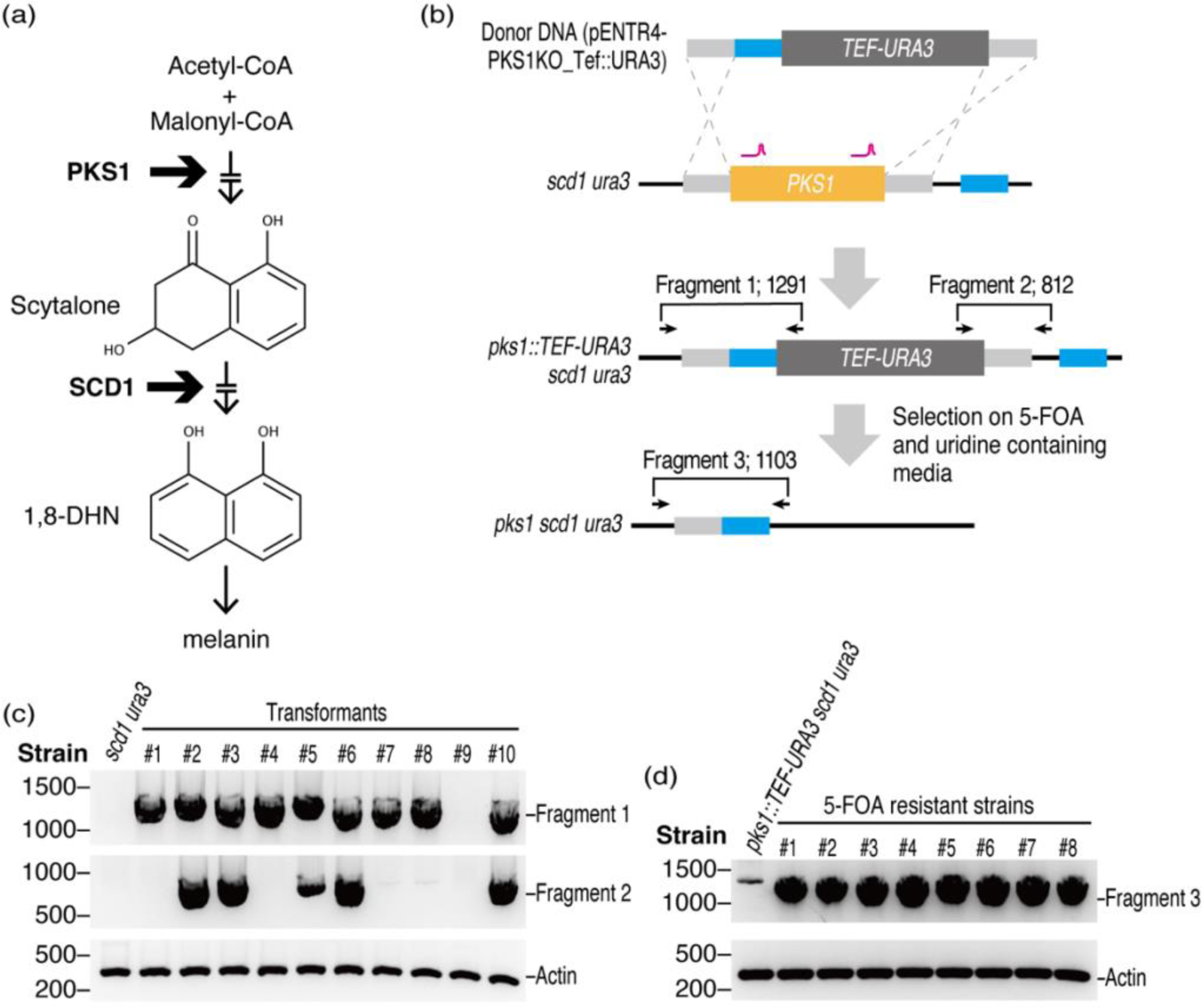
*URA3*-based marker recycling systems is applicable for *PKS1* knockout in *scd1 ura3.* (a) Schematic of assumed melanin biosynthesis pathway in *C. higginsianum*. Thin arrows indicate the flow of the melanin biosynthesis, and thick arrows indicate the modifications of melanin precursors by PKS1 or SCD1. Gaps inside the arrows indicate that several melanin precursors are omitted for simplified understanding. (b) Construction design of *PKS1* knock-out and the removal of *TEF-URA3*. The donor DNA has *TEF-URA3*, flanked by 0.5 kb homology arms depicted as light gray boxes. Lines in magenta represent the sites targeted by gRNAs. Light blue boxes show homology arms to remove *TEF-URA3* from the genome of *pks1::TEF-URA3 scd1 ura3* via homologous recombination. Arrows represent primers used to amplify “Fragment 1” and “Fragment 2” existing specifically in the genome of *pks1::TEF-URA3 scd1 ura3* and “Fragment 3” in the genome of *pks1 scd1 ura3*. (c) PCR screening of *pks1::TEF-URA3 scd1 ura3* mutants. PCRs were performed using the primer sets depicted in (b) (Fragment 1 and 2). Results from randomly selected 10 colonies from transformants are shown. (d) PCR screening of *pks1 scd1 ura3* mutants. PCRs were performed using primer sets depicted in (b) (Fragment 3). Results from randomly selected eight colonies from transformants are shown.

### 2.4 *URA3* knock-in complements the uridine auxotrophy of *ura3* mutants

To complement the uridine auxotrophy of *ura3* mutants, we knocked-in *C. higginsianum URA3* to *ura3* mutants utilising the CRISPR-Cas9 system. The donor DNA for *URA3* knock-in contained an open reading frame of *URA3* with two homology arms that are homologous to upstream and downstream of the *NPTII* expression cassette (Figure 5a). Two gRNAs targeting the *NPTII* expression cassette were designed and the transformation of *ura3* using the donor DNA and RNPs resulted in the generation of more than 120 colonies. The PCR screening of 50 randomly-selected colonies revealed that 96% of colonies (48 colonies) were *URA3* knocked-in strains (*URA3* KI) (Figure 5b).

**Figure 5.**
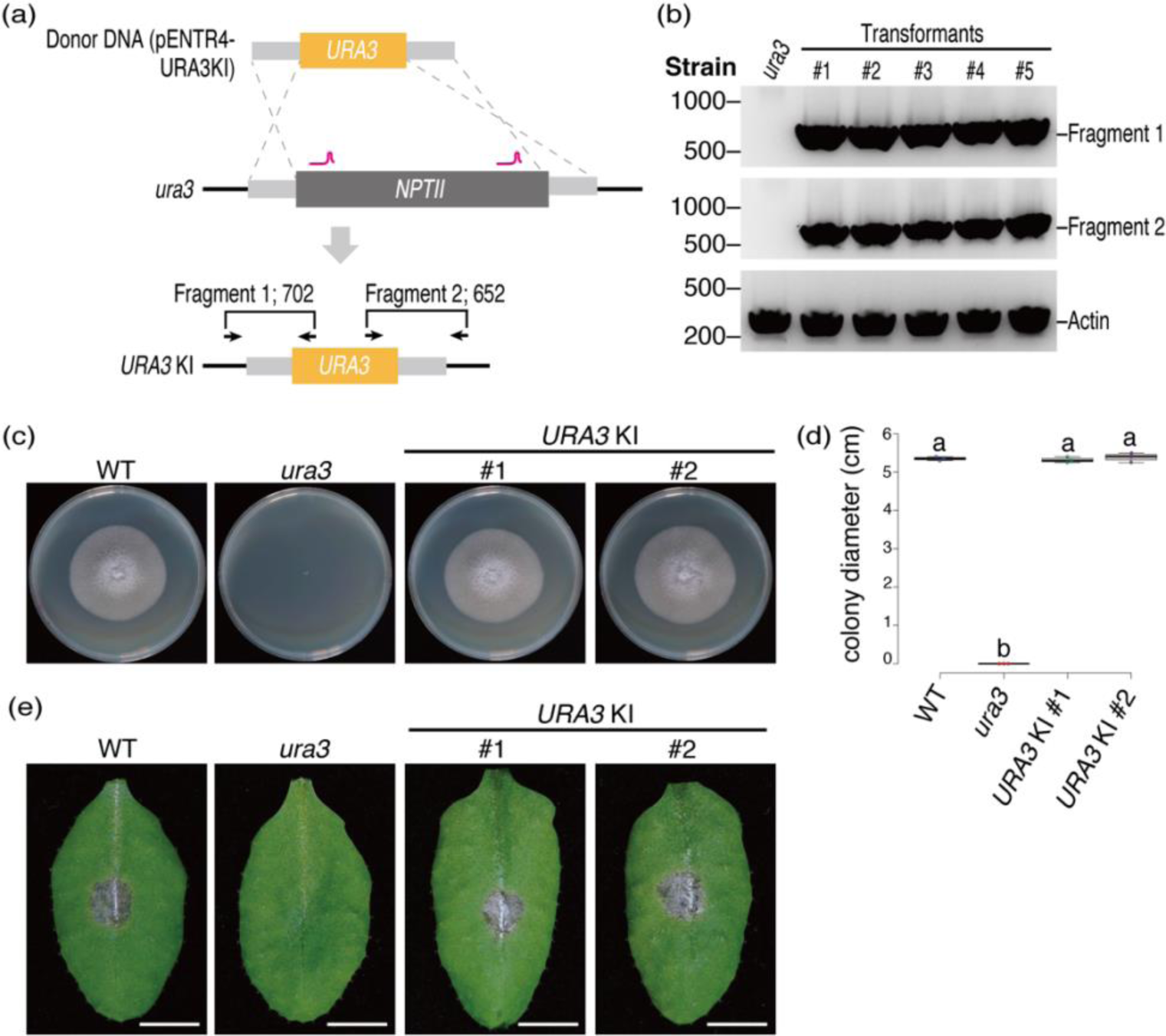
*URA3* knock-in to *ura3* mutants complements the uridine auxotrophy. (a) Construction design of *URA3* knock-in to *ura3* mutants. The donor DNA has the *URA3* gene, flanked by 0.5 kb homology arms depicted as light gray boxes. Lines in magenta represent the sites targeted by gRNAs. Arrows represent primers used to amplify “Fragment 1” and “Fragment 2” existing specifically in the genome of *URA3* KI. (b) PCR screening of *URA3* KIs. PCRs were performed using the primer sets depicted in (a). Results from randomly selected five colonies from transformants are shown. (c) Normal colony growth of two independent *URA3* KI strains on MA. Photographs were taken at seven days post-culture. (d) Box plots of colony diameters of *C. higginsianum* strains shown in (c). Different characters in each graph indicate statistical significance (Tukey HSD, *n*=3). (e) Disease symptoms on *A. thaliana* leaves after inoculation with each genotype of *C. higginsianum*. Leaves were observed at seven days post-inoculation. The brownish lesion observed after inoculation with wild-type *C. higginsianum* and *URA3* KIs was not observed after inoculation with *ura3* mutants (*n*=6). Bars represent 5 mm.

To confirm the complementation of uridine auxotrophy of *URA3* KI, we measured the colony diameters on media. Colonies of *URA3* KIs were as large as those of the wild type (Figure 5c, d), indicating that the uridine auxotrophy of *ura3* mutants was complemented by *URA3* knock-in. In addition, we assessed the requirement of *URA3* for the pathogenicity of *C. higginsianum* to *A. thaliana*. The inoculation of wild-type *C. higginsianum* caused brownish anthracnose disease symptoms, whereas no symptoms were observed in the *ura3* mutant. The symptomless phenotype of *ura3* was restored by *URA3* knock-in (Figure 5e), indicating the requirement of *URA3* for the pathogenicity of *C. higginsianum*. Therefore, our results show the successful complementation of uridine auxotrophy and pathogenicity by *URA3* knock-in using the CRISPR-Cas9 system in *C. higginsianum*.

### 2.5 *scd1* and *pks1 scd1* show melanin-deficient phenotypes

Given that melanin plays a role in pathogenicity by conferring physical rigidity to appressoria of certain phytopathogenic fungi (Kubo et al., 1991; Ludwig et al., 2014; Woloshuk et al., 1980), we investigated the melanisation of *scd1* and *pks1 scd1* mutants. We first complemented the uridine auxotrophy of *scd1 ura3* and *pks1 scd1 ura3* mutants by knocking-in the *URA3* gene (Figure S2). Subsequently, we examined the melanisation of *in vitro* appressoria of wild-type *C. higginsianum, scd1* and *pks1 scd1* mutants under a microscope. While the appressoria of wild-type *C. higginsianum* were black, those of *scd1* and *pks1 scd1* mutants were transparent (Figure 6a), indicating the lack of melanin. We then evaluated the pathogenicity of *scd1* and *pks1 scd1* mutants on *A. thaliana* leaves. Notably, no symptoms were observed after inoculation with *scd1* or *pks1 scd1* mutants (Figure 6b), suggesting that melanin is required for the pathogenicity of *C. higginsianum* on *A. thaliana*.

**Figure 6.**
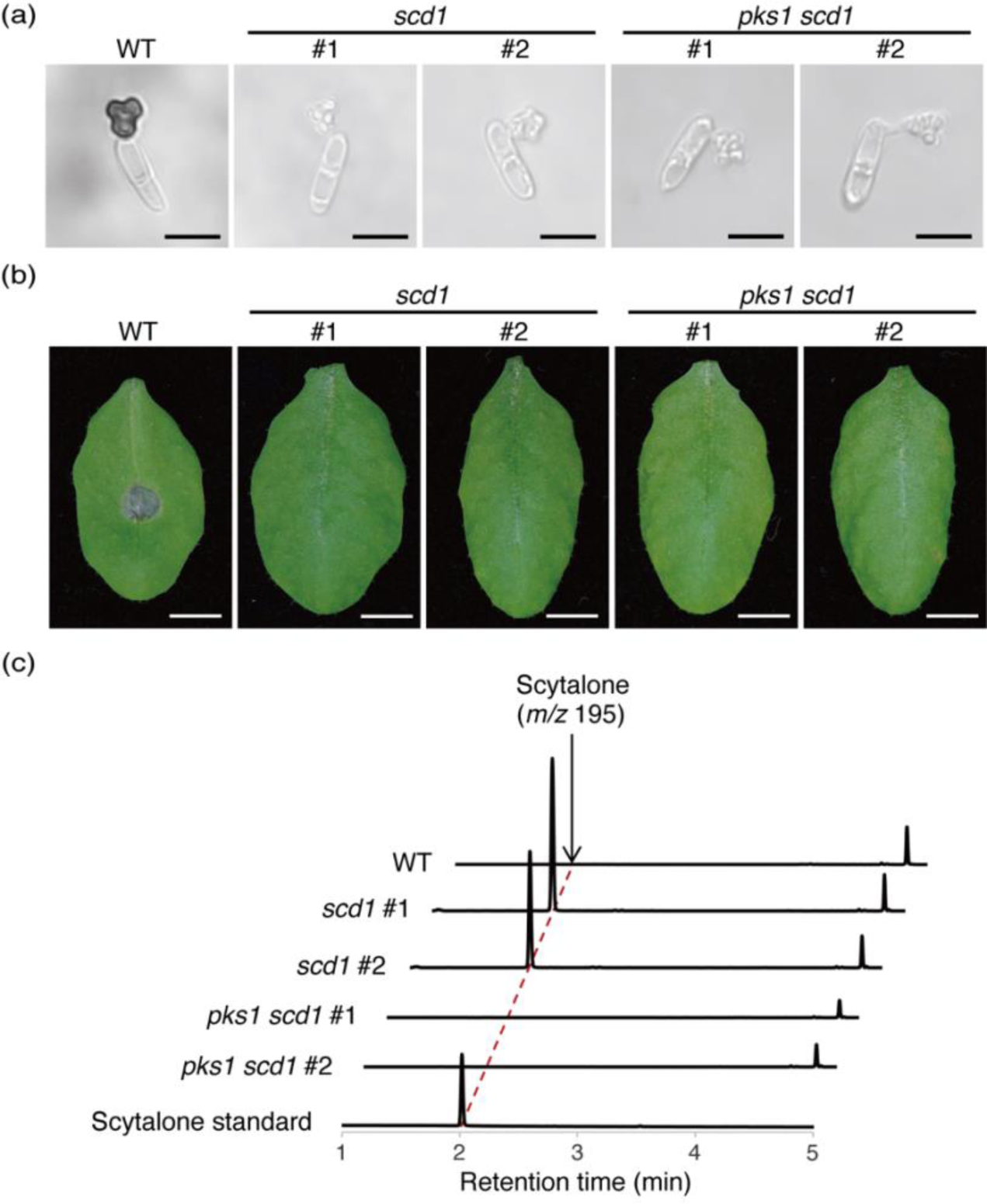
Melanin-deficient phenotypes of *scd1* and *pks1 scd1* mutants. (a) Deficiency of melanisation in *in vitro* appressoria of *scd1* and *pks1 scd1* strains. Appressoria were observed after incubating conidia on glass bottom dishes in a humid box at 25°C for 18-19 hours. Bars represent 10 µm. (b) Loss of disease symptoms by *scd1* and *pks1 scd1* inoculation. Leaves were observed at seven days post-inoculation (*n*=6). Bars represent 5 mm. (c) Accumulation of scytalone in appressoria of the *scd1* strain. Metabolites were extracted from *in vitro* appressorial cells of *C. higginsianum* strains and analysed by UPLC/MS. Metabolites were detected by absorbance at 281 nm.

We compared the accumulation of scytalone, a metabolite assumed to be a substrate of SCD1, by exploiting the established method for mass-producing *in vitro* appressoria of *C. higginsianum* (Kleemann et al., 2008). We expected that scytalone would accumulate only in *scd1* but not in wild-type *C. higginsianum* or *pks1 scd1* mutants, since PKS1 and SCD1 were assumed to produce a precursor of scytalone and dehydrate scytalone, respectively (Takano et al., 1997). To detect scytalone, we used ultra-performance liquid chromatography (UPLC)/mass spectrometry (MS) to analyse metabolites from *in vitro-* induced *C. higginsianum* appressoria. As expected, scytalone was detected only in *scd1* mutants, but not in wild-type *C. higginsianum* or *pks1 scd1* mutants (Figure 6c). These results support the idea that PKS1 and SCD1 play roles in the upstream and downstream steps of scytalone synthesis in the melanin biosynthetic pathway, respectively (Figure 4a). Moreover, the successful double knock-out of *scd1* and *pks1* in the same strain was confirmed, as indicted by our results.

## 3 DISCUSSION

The challenges posed by the low efficiency of gene disruption and redundancy of virulence factors have hindered the search for virulence factors in many phytopathogenic fungi, including *C. higginsianum* (Collemare et al., 2019; Ushimaru et al., 2010). In this study, we addressed these challenges by introducing two methods, the CRISPR-Cas9 system and the *URA3*-based marker recycling, thereby establishing a highly efficient method for multiple gene disruption in *C. higginsianum*. Our results demonstrated that CRISPR-Cas9 significantly increased the efficiency of gene disruption by donor DNA-mediated homologous recombination (Figures 1c and 3b). Moreover, we confirmed the feasibility of the *URA3*-based marker recycling approach in *C. higginsianum*, as evidenced by the presence of a single *URA3* homologue and the demonstration of uridine auxotrophy and 5-FOA insensitivity in the *ura3* mutant (Figure 2). Using these methods, we successfully disrupted two melanin biosynthetic genes, *SCD1* and *PKS1*, and confirmed that the sequential knock-out of these genes was possible with high efficiency (Figures 3c, e and 4c, d). Phenotypic analyses revealed that *scd1* and *pks1 scd1* mutants failed to melanise and lost virulence (Figure 6), thereby confirming the effectiveness of our method for analysing virulence factors. Overall, our new method is a significant addition to the toolkit for *C. higginsianum* reverse genetics and will facilitate the future identification of virulence factors, which was previously challenging.

Two methods can be employed for CRISPR-Cas9-based fungal transformation: direct application of *in vitro* formed RNPs and DNA-mediated expression of RNPs (Schuster and Kahmann, 2019). In this study, we utilised the *in vitro* formed RNP approach (Figure 1a), which presents two benefits and one drawback. Firstly, the use of *in vitro*-formed RNPs can minimise off-target effects and cytotoxicity as RNPs are present in the cell for a limited time. DNA-mediated expression of RNPs runs the risk of Cas9 and gRNA expression cassettes being integrated into the genome and persistently expressed post-genome editing, leading to off-target effects or cytotoxicity (Foster et al., 2018; Jacobs et al., 2014; Zhang et al., 2015). Secondly, there is no need to scrutinise Cas9 and gRNA expression promoter conditions. Selecting appropriate promoters and codon optimisation are generally necessary for DNA-mediated approaches to induce DSBs (Morio et al., 2020; Nødvig et al., 2015). RNP-based methods do not require these efforts, and determining the required amount of Cas9 and gRNAs for genome editing is straightforward. The disadvantage of the RNP-based approach is its current costliness. DNA-mediated approaches are less expensive than RNP-based approaches using commercially available recombinant Cas9 proteins and synthetic gRNAs since they only require DNA constructs. However, this disadvantage can be remedied by preparing the Cas9 protein in-house instead of acquiring it commercially. In our study, a single purification yielded approximately 10 mg of Cas9 protein, more than 100 times the amount necessary for *C. higginsianum* transformation.

The application of CRISPR-Cas9 for improving homologous recombination efficiency offers significant advantages over traditional methods suppressing NHEJ. The conventional approaches that target key NHEJ factors such as *Ku70* or *Lig4* homologues have shown promise in reducing the frequency of the NHEJ-mediated DSB repair and thereby increasing the HDR-mediated homologous recombination frequency (Ninomiya et al., 2004; Ushimaru et al., 2010; Zhang et al., 2021). However, this approach has the drawback of disrupting the innate eukaryotic DNA repair mechanism, potentially leading to genome instability and unexpected homologous recombination events (Pastink et al., 2001; Shrivastav et al., 2008). In contrast, the CRISPR-Cas9 method is genetically cleaner, as it does not require the disruption of *Ku70* or *Lig4* homologues. Furthermore, CRISPR-Cas9 can enhance transformation efficiency by inducing DSBs repaired with donor DNAs (Figure 1d). Apart from these advantages, CRISPR-Cas9 has broad applicability beyond homologous recombination.

The CRISPR-Cas9 system can induce mutations without donor DNAs when combined with NHEJ. In *B. cinerea* and *M. oryzae*, CRISPR-Cas9-induced DSBs have been shown to mutate target genes, with NHEJ repairing the break and introducing a deletion or insertion of several bases (Foster et al., 2018; Leisen et al., 2020). Since NHEJ cannot introduce selection markers, unlike donor DNA-mediated homologous recombination, selecting transformants is challenging, in theory. However, in *B. cinerea* and *M. oryzae*, this issue was tackled by co-transforming donor DNAs with selection markers along with CRISPR-Cas9 RNPs and isolating them on a selective medium, which is called “co-editing”. This method’s key advantage is that it can simultaneously disrupt multiple genes by using several guide RNAs. Nevertheless, the lack of selection markers for each gene requires extensive screening efforts to isolate the desired multiple mutants when the transformation efficiency is not as high as in *B. cinerea*. If transformation efficiency can be further improved in *C. higginsianum*, this method is an attractive technique to apply.

Compared to other marker recycling systems such as Cre-*loxP* and Flp-*FRT*, the *URA3*-based system has two notable advantages and one disadvantage. The first advantage is that the *URA3*-based marker recycling process does not leave any foreign sequences on the genome, whereas the Cre-and Flp-based systems excise DNA between two recombination sites, respectively, leaving one site in the recipient genome. In contrast, the integrated foreign sequence can be entirely removed through homologous recombination in the *URA3*-based system, resulting in cleaner knock-out lines (Figures 3d and 4b). The second advantage is its simple donor DNA construction process. Cre-*loxP* requires seven sequences for donor DNA: three gene expression cassettes (positive and negative selection markers and a *Cre* gene expression cassette), two flanking *loxP* sequences and two flanking homology arms. In contrast, the *URA3*-based system requires only four sequences: an *URA3* expression cassette and three flanking homology arms, making it easier to construct. However, the *URA3*-based system has a disadvantage, unlike Cre-*loxP* and Flp-*FRT*, that the parental strain requires an *URA3* mutation. The *URA3* mutation leads to a loss of virulence phenotype, which necessitates the knock-in of *URA3* to the *ura3* background when assessing virulence on plants (Figure 5c). Nonetheless, this drawback can be overcome by using highly efficient *URA3* knock-in methods such as CRISPR-Cas9, with the *URA3* knock-in rate reaching 96% (Figure 5b).

In conclusion, this study presents a highly efficient method for disrupting multiple genes in *C. higginsianum*, including genes that were previously challenging to target. In principle, our method can be applied to fungi capable of transforming protoplasts and carrying *URA3* homologs (Figure S1). Indeed, we have previously utilised *URA3*-based marker recycling (Kumakura et al., 2019) and CRISPR-Cas9 (Chen et al., 2023) in *C. orbiculare*. The method we developed has a lower throughput compared to methods reported in organisms that introduce multiple mutations simultaneously by combining RNP-induced DSBs and NHEJ (Leisen et al., 2020; Stuttmann et al., 2021). However, we have demonstrated the potential to combine our *in vitro*-formed RNPs with NHEJ to perform multiple mutageneses simultaneously in *C. higginsianum*. With further improvements in RNP introduction efficiency, this approach can be utilised for easier and faster disruption of multiple target genes.

## 4 EXPERIMENTAL PROCEDURES

### 4.1 Accession numbers

Accession numbers of *C. higginsianum* genes targeted in this study, *URA3*, *SCD1* and *PKS1*, are listed in Table S1.

### 4.2 Plasmid constructions

*C. higginsianum* genomic DNA used as PCR templates were isolated by Maxwell RSC Plant DNA Kit with Maxwell RSC 48 instrument (Promega) following the manufacturer’s protocol. PCRs were performed using KOD One PCR Master Mix −Blue-(Toyobo) DNA polymerases. All plasmids used in this study are listed with their contents in Table S2, and primers to construct plasmids are listed with their sequences in Table S3. Briefly, fragments for fungal transformation were fused into pENTR4 plasmids (Thermo Fisher Scientific) by HiFi DNA assembly (New England Biolabs). *NPTII* expression cassette was amplified using pII99 plasmid (Namiki et al., 2001).

### 4.3 CRISPR-Cas9 RNP preparation for transformation

#### 4.3.1 Cas9 recombinant protein preparation

Cas9 recombinant protein fused with simian virus 40 nuclear localisation signal and his-tag at C-terminus (Cas9-NLS-His) was expressed in *E. coli* strain Rosetta 2 (DE3) cells (Novagen) transformed with pET-Cas9-NLS-6xHis (Addgene plasmid #62933) (Zuris et al., 2015). Cas9-NLS-His protein expression was induced by overnight incubation at 20°C with isopropyl β-D-1-thiogalactopyranoside at a final concentration of 0.4 mM. The harvested cells were resuspended in immobilised metal chromatography (IMAC) buffer (50 mM Tris-HCl [pH 7.5], 300 mM NaCl) supplemented with 3 kU/ml rLysozyme (Novagen). The cell suspension was sonicated on ice (Ultrasonic disruptor UD-201, TOMY), and the insoluble debris were removed by centrifugation twice (10,000 *g*, 10 minutes, 4°C). The supernatant was applied to an IMAC column (HisTALON superflow cartridge, Clontech) pre-equilibrated with IMAC buffer. The column was washed with IMAC buffer supplemented with 5 mM Imidazole, and the bound protein was eluted with IMAC buffer supplemented with 150 mM Imidazole. The eluted fractions containing Cas9-NLS-His were diluted 4-fold with 50 mM Tris-HCl (pH 7.5) and subjected to cation exchange chromatography (Hitrap S HP, GE healthcare) pre-equilibrated with 50 mM Tris-HCl (pH 7.5) and 100 mM NaCl. The bound proteins were eluted in a linear NaCl gradient (100 – 750 mM). The fractions containing Cas9-NLS-His were pooled and buffer-exchanged into 10 mM Tris-HCl (pH 7.5), 300 mM NaCl, 0.1 mM EDTA, 1 mM DTT, 40% Glycerol. This Cas9 recombinant protein was stored at −20°C until use.

#### 4.3.2 gRNA design and preparation

Targeting sequences of gRNAs were determined through the following two steps: Candidate sequences were predicted from the target regions with target cleavage efficacy by the web-based CRISPick software (Doench et al., 2016; Sanson et al., 2018): For each candidate sequence, the likelihood of off-targeting in the *C. higginsianum* genome (GenBank accession: PRJNA47061) was calculated using CasOT (Xiao et al., 2014), and those with lower off-targeting likelihood and higher target cleavage efficiency were selected. Then, gRNAs were synthesised by GenScript or IDT. To enhance the chances of target cleavage, two distinct gRNAs were synthesised for each target gene. Target sequences of gRNA are listed in Table S4.

#### 4.3.3 CRISPR-Cas9 RNP preparation

To prepare RNPs, gRNA was incubated at 95°C for three minutes, cooled on ice, mixed with Cas9 recombinant protein, and incubated for five minutes at room temperature. For each transformation, RNPs (∼400 pmol) consisting of Cas9 recombinant protein and gRNAs were used.

### 4.4 *C. higginsianum* transformation

The parental strain used was *C. higginsianum* (Strain ID: IMI349063A) (Dallery et al., 2017). All transformation experiments were performed under sterile conditions. The transformation of *C. higginsianum* strains with *URA3* mutations was performed in the same way as that of *C. orbiculare*, as reported in previous reports (Chen et al., 2023; Kumakura et al., 2019) with modifications: *C. higginsianum* strains with *URA3* mutation were cultured on Mathur’s media agar (MA) (16 mM glucose, 5 mM MgSO_4_, 20 mM KH_2_PO_4_, OXOID mycological peptone 0.22% [w/v], agar 3% [w/v]) (Mathur, R.S., Barnett, H.L., & Lilly, V.G., 1950) containing 10 mM uridine (Fujifilm Wako), and 0.6 M glucose minimal media (1.6 g/l yeast nitrogen base w/o amino acids [Difco], 4 g/l L-Asparagine-Monohydrate, 1 g/l NH_4_NO_3_, 3% [w/v] Bacto agar, 0.6 M glucose, pH6.0) was used for the selection of the *URA3* coding sequence containing transformants. The transformation of *C. higginsianum* strains without *URA3* mutation was performed as follows.

#### 4.4.1 Conidial preparation

Glycerol stocks of *C. higginsianum* strains stored at −80°C were streaked on MA in 90 mm dish and incubated at 25°C in the dark. After seven to nine days, pieces of agar with fungal cells were transferred onto 100 ml MA in four to five 250 ml flasks by sterilised plastic straws, 1 ml sterilised water was added and mixed well to spread fungal cells on the entire surface of the MA plate. Flasks were covered with aluminium foil and sponge plugs for aeration and incubated at 25°C in the dark for seven to ten days.

#### 4.4.2 Protoplast preparation

Conidia in flasks were suspended by Mathur’s liquid media (16 mM glucose, 5 mM MgSO_4_, 20 mM KH_2_PO_4_, OXOID mycological peptone 0.22% [w/v]) (7.5 ml per flask) and filtered through a cell strainer (100 µm pore size, Corning). Then, the conidia suspension in 100 ml Mathur’s liquid media (∼8×10^7^ conidia) was incubated in a sterilised 300 ml flask for germ tube induction (25°C, 100 rpm, 14-17 hours, rotary shaker). Four to five flasks were cultured for each protoplast preparation. Cells were harvested using a cell strainer (100 µm pore size), washed with 0.7 M NaCl, and resuspended in a 20 ml filter sterilised (0.2 µm pore size, MERK) digestion mix (1.2 M MgSO_4_, 10 mM NaPO_4_, 3% [w/v] lysing enzyme from *Trichoderma harzianum* [MERK], pH5.6). After gently agitating for three to four hours (60 rpm, 25°C, rotary shaker), the suspension was then underlaid with 20 ml of trapping buffer (0.6 M sorbitol, 10 mM Tris-HCl pH7.5, 50 mM CaCl_2_) and subsequently centrifuged (960 *g*, five minutes, swinging-bucket rotor). The protoplasts were isolated from the interface between the two layers, rinsed twice with STC (1.2 M sorbitol, 10 mM Tris-HCl pH7.5, 50 mM CaCl_2_), and re-suspended in STC (10^7^-10^8^ protoplast ml^−1^). 100-150 µl aliquots were stored at −80°C.

#### 4.4.3 Transformation

Thawed protoplasts were mixed with donor DNA (∼10 µg) and CRISPR-Cas9 RNPs (∼400 pmol) and incubated on ice for 10-20 minutes. PEG solution (60% [w/v] PEG4000, 10 mM Tris-HCl pH7.5, 50 mM CaCl_2_) was added in three steps (200 µl, 200 µl, and 800 µl) with five minutes incubation on ice after each addition. After adding 1 ml of STC, protoplasts were mixed with 200 ml of regeneration medium (0.1% [w/v] yeast extract, 0.1% [w/v] casein acid hydrolysate [MERK], 1 M sucrose, 2.4% [w/v] Bacto agar) and spread in square plastic Petri dishes (140×100 mm, 40 ml per plate). Dishes were incubated for six to sixteen hours, then overlaid with 20 ml of top media (3% [w/v] Bacto agar) containing 500 µg/ml G418 (Fujifilm Wako) and incubated at 25°C for five to seven days. The resulting colonies were transferred to MA containing 500 µg/ml G418 and incubated for three to five days. Transformants were identified as colonies that grew on this second selective medium.

### 4.5 Selection of the strains without *TEF-URA3*

*TEF-URA3* cassettes were removed from *TEF-URA3* containing strains basically as previously reported (Kumakura et al., 2019). Briefly, 2 to 5×10^5^ conidia were spread on MA containing 1 mg/ml 5-FOA (Fujifilm Wako) and 10 mM uridine and incubated.

### 4.6 PCR screening and storage

The genomic DNA of each transformant was isolated following a previously reported method (Liu et al., 2000) and subjected to PCR to confirm the desired transformation. All derivative fungal strains and DNAs used for transformation in this study are listed in Table S5 and Table S2, respectively. Primers used for PCR screening are listed in Table S6. Conidia of selected strains were mixed with glycerol at 25% of the final concentration and stored at −80°C for long-term storage.

### 4.7 C. higginsianum inoculation on A. thaliana

Five-week-old leaves of *Arabidopsis thaliana* Col-0 grown at 22°C under an 8 hr light/16 hr dark cycle were used for inoculation. Glycerol stocks of *C. higginsianum* strains stored at −80°C were streaked on MA and incubated at 25°C in the dark. After seven to nine days, pieces of agar with fungal cells were transferred onto 100 ml MA in a 250 ml flask, added 1 ml sterilised water, mixed well. Flasks were covered with aluminium foil and sponge plugs for aeration and incubated at 25°C in the dark. After seven to nine days, conidia were suspended in sterilised water and concentration was measured by a hemacytometer. Ten µl of the conidial suspensions (1×10^6^ conidia ml^−1^) were drop-inoculated onto *A. thaliana* leaves and incubated under the same condition as the plant growth in a humid box. After seven days, the inoculated leaves were detached from plants and photographed.

### 4.8 *In vitro* appressoria preparation

#### 4.8.1 Conidia preparation

Conidia were prepared as described in the “conidial preparation” section, with a final concentration of 2×10^6^ conidia ml^−1^.

#### 4.8.2 For microscopic observation

The conidial suspension was applied to a 35 mm glass bottom dish (ibidi, IB81218) using a disposable hand-held plastic spray. The inoculated dishes were incubated at 25°C in a humid box for 18-19 hours. Appressoria were observed and photographed using the BX51 microscopy (Olympus) and the LP74 microscope digital camera (Olympus). The UPlanSApo 40× objective lens (Olympus) was used.

#### 4.8.3 For appressoria lysate

To produce large quantities of *in vitro* appressoria, a modified version of a previously reported method was used (Kleemann et al., 2008). Briefly, 40 ml of conidia suspension in sterilised distilled water (2×10^6^ conidia ml^−1^) was added to a polystyrene dish (14×10 cm) and allowed to settle and adhere to the surface for 40 minutes. A sterilised nylon mesh (13×9 cm, 50 µm pore size) was placed on the surface of the liquid and the excess water was discarded, leaving the nylon mesh to retain water on the dish. After 18-19 hours incubation at 25°C in a humid box, appressoria formed on the bottom of polystyrene dish was harvested with cell scraper together with 1 ml sterilised distilled water. The collected fluid was transferred to 2 ml steel-top tubes and homogenised using 0.6 mm and 5 mm zirconia beads (BMS) in a ShakeMaster Neo (BMS) at 2,000 rpm for two minutes. The fluid containing crushed appressoria was collected and stored at −80°C for further analysis.

### 4.9 Metabolite analysis

Appressoria crushed fluid was treated with four volumes of ethanol, and the supernatant was dried under nitrogen. The dried material was dissolved in methanol and analysed using UPLC/MS on a Waters Acquity UPLC H-Class-QDa system (Waters, Milford, MA). A reversed-phase column (BEH C18, 2.1×50 mm, 1.7 µm, Waters) was used at a flow rate of 0.6 ml min^−1^. The gradient system was acetonitrile (solvent B) in 0.1% formic acid-water (solvent A): 5% B from 0 to 1 minute, 5% to 95% B from 1 to 4.5 minutes, 95% B from 4.5 to 5.5 minutes, and re-equilibration with 5% B from 5.5 to 8 minutes. Ultraviolet absorption and positive ion electrospray ionisation were used for detection of metabolites. Scytalone was prepared from the cultured broth of carpropamid-treated *Pyricularia oryzae* P2 strain as described previously (Kurahashi et al., 1998).

## 5 ACKNOWLEDGEMENTS

This work was supported by RIKEN Junior Research Associate Program to K.Y, JSPS (23K05158, JP20K15500, 19KK0397 and JP18K14440) and Japan Science and Technology Agency (JST) (ACT-X, JPMJAX20B4) to N.K., and JSPS (JP20H05909 and JP22H00364) to K.S. We thank Akiko Ueno, Takuma Aita, Hayato Fukuda, and Reika Hiraishi for their technical assistance.

## 6 DATA AVAILABILITY STATEMENT

Data sharing is not applicable to this article as no new data were created or analysed.

