## Supplementary material for "Efficient multiple gene knock-out in *Colletotrichum higginsianum* via CRISPR-Cas9 ribonucleoprotein and *URA3*-based marker recycling": Figure S1

| Phylum | Class | | # of *URA3* homologues |
|---|---|---|---|
| B | 1 | *Puccinia graminis* | 1 |
| | 2 | *Ustilago maydis* | 1 |
| | 3 | *Melampsora lini* | - |
| Ascomycota | 4 | *Saccharomyces cerevisiae* | 1 |
| | | *Candida albicans* | 1 |
| | 5 | *Zymoseptoria tritici* | 1 |
| | 6 | *Blumeria graminis* | 1 |
| | | *Botrytis cinerea* | 1 |
| | Sordariomycetes | *Neurospora crassa* | 1 |
| | | *Pyricularia oryzae* | 1 |
| | | *Fusarium graminearum* | 1 |
| | | *Fusarium oxysporum* | 1 |
| | | *Colletotrichum orbiculare* | 2 |
| | | ***Colletotrichum higginsianum*** | 1 |

B: Basidiomycota
1: Teliomycetes, 2: Ustilaginomycetes
3: Urediniomycetes, 4: Saccharomycetes
5: Dothideomyetes, 6: Leotiomycetes

**Figure S1** Number of *URA3* genes in selected fungi including *C. higginsianum*. Homologues of *URA3* genes were predicted by a NCBI BlastP search in the default setting using *S. cerevisiae* Ura3p as a query. Species in green letters are phytopathogenic fungi.
