## Supplementary material for "Efficient multiple gene knock-out in *Colletotrichum higginsianum* via CRISPR-Cas9 ribonucleoprotein and *URA3*-based marker recycling": Figure S2

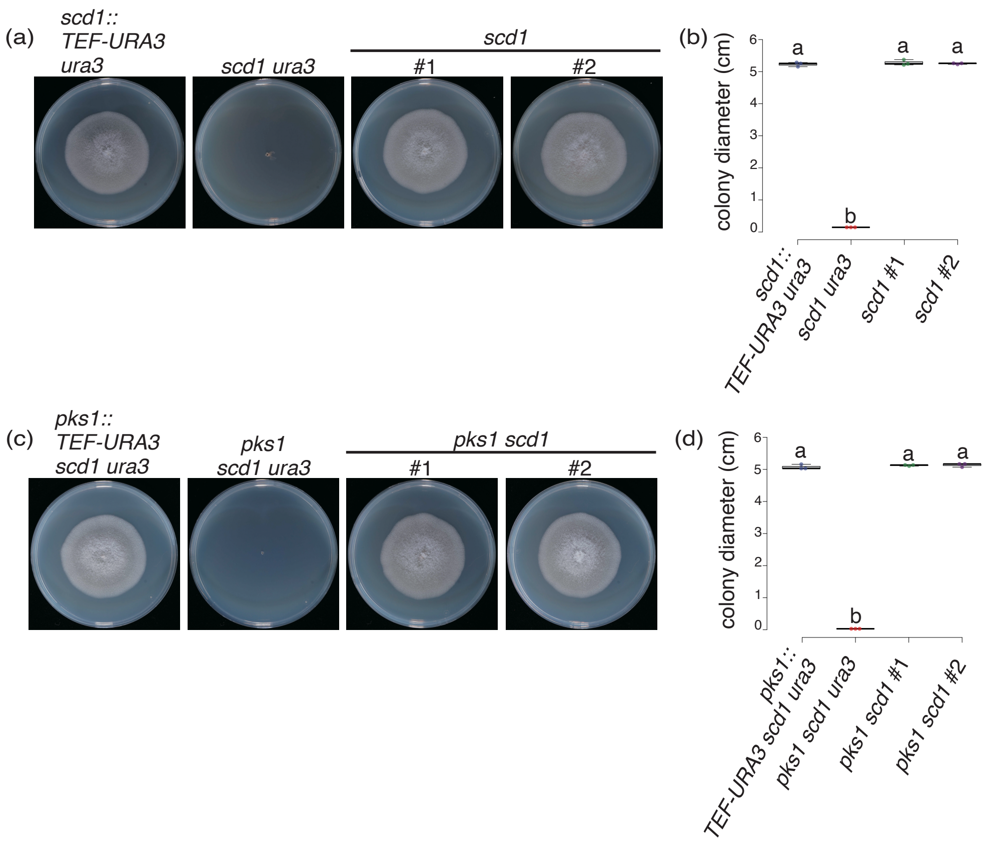


**Figure S2** Complementation of uridine auxotrophy of *scd1 ura3* and *pks1 scd1 ura3* strains by *URA3* knock-in.

(a) Normal colony growth of two independent *scd1* strains on MA. Photographs were taken at seven days post-culture (dpc).

(b) Box plots of colony diameters of *C. higginsianum* strains shown in (a). Different letters in each graph indicate statistical significance (Tukey HSD, *n*=3).

(c) Normal colony growth of two independent *pks1 scd1* strains on MA. Photographs were taken at 7 dpc.

(d) Box plots of colony diameters of *C. higginsianum* strains shown in (c). Different letters in each graph indicate statistical significance (Tukey HSD, *n*=3).
